# Helicity of a tardigrade disordered protein promotes desiccation tolerance

**DOI:** 10.1101/2023.07.06.548010

**Authors:** Sourav Biswas, Edith Gollub, Feng Yu, Garrett Ginell, Alex Holehouse, Shahar Sukenik, Thomas C. Boothby

## Abstract

In order to survive extreme drying (anhydrobiosis), many organisms, spanning every kingdom of life, accumulate intrinsically disordered proteins (IDPs). For decades, the ability of anhydrobiosis-related IDPs to form transient amphipathic helices has been suggested to be important for promoting desiccation tolerance. However, evidence empirically supporting the necessity and/or sufficiency of helicity in mediating anhydrobiosis is lacking. Here we demonstrate that the linker region of CAHS D, a desiccation-related IDP from tardigrades that contains significant helical structure, is the protective portion of this protein. Perturbing the sequence composition and grammar of the linker region of CAHS D, through the insertion of helix-breaking prolines, modulating the identity of charged residues, sequence scrambling, or replacement of hydrophobic amino acids with serine or glycine residues results in variants with different degrees of helical structure. Importantly, the resulting helicity of these variants generated through similar helix breaking modalities correlates strongly with their ability to promote desiccation tolerance, providing direct evidence that helical structure is necessary for robust protection conferred by this desiccation-related IDP. However, correlation of protective capacity and helical content in variants generated through different helix perturbing modalities do not show as strong a trend, suggesting that while helicity is important it is not the only property that makes a protein protective during desiccation. These results provide direct evidence for the decades old theory that helicity of desiccation-related IDPs is linked to their anhydrobiotic capacity.

## Introduction

Water is required for all metabolism, so the adage “water is life” is commonly used to imply that without water, life cannot persist^1^. However, diverse organisms spanning every kingdom of life contradict this statement through their ability to enter into a drying-induced state of suspended animation known as anhydrobiosis (“life without water”)^2^. In this anhydrobiotic state, these organisms can persist for years, decades, in the case of some ancient seeds, even millennia, but resume metabolism and life processes upon rehydration^3–5^.

Decades of work have identified non-reducing disaccharides, such as trehalose, as major molecular mediators that promote anhydrobiosis in a number of organisms^6–8^. However, more recent work has demonstrated that some organisms that robustly survive desiccation, such as tardigrades and bdelloid rotifers, produce low or undetectable levels of trehalose^9–18^. While these discoveries do not reduce the importance or necessity of trehalose and other co-solutes in mediating desiccation tolerance in some organisms, they do reveal that additional mediators likely play important roles in surviving desiccation.

An emerging paradigm in the anhydrobiosis field is the involvement of intrinsically disordered proteins (IDPs) in mediating desiccation tolerance^16, 19, 20^. Some of the first IDPs implicated in anhydrobiosis were late embryogenesis abundant (LEA) proteins, which were first identified in cotton seeds^21^, but have subsequently been found in a number of diverse organisms^17, 21–26^. Since their discovery, LEA proteins have been classified into seven distinct families, and other desiccation-related IDPs have been identified^27–30^. These include LEA-like proteins or hydrophilins^27^, which are typified by a predominance of hydrophilic residues^31^, as well as three families of tardigrade disordered proteins (TDPs) termed cytoplasmic, mitochondrial, and secreted abundant heat soluble (CAHS, MAHS, and SAHS, respectively) proteins^32, 33^.

While IDPs lack a stable tertiary structure, this does not preclude them from adopting transient secondary structure^34^. Members of several LEA families, as well as CAHS, MAHS, SAHS proteins, have been shown to possess transient helices and/or to take on additional helicity during drying and/or chemical desolvation^26, 28, 30, 35–37^. Adoption of helical structure in these proteins often results in the appearance of amphipathicity, that is, the appearance of a hydrophobic and polar face^28, 38^. This phenomenon has been described in a myriad of studies, many of which propose that helicity of these proteins drives their protective capacity during anhydrobiosis^28, 36, 39^.

Although helicity has been proposed to play a role in desiccation protection, direct evidence testing this hypothesis is limited. One study identified a correlation between freeze tolerance and helicity in a cold-regulated LEA protein^40^. However, to our knowledge, no published work has directly tested the link between the helicity of a desiccation-related IDP and its ability to confer protection during drying.

Here we directly test the necessity of the helicity of the tardigrade IDP, CAHS D, in protecting the labile enzyme lactate dehydrogenase (LDH) during desiccation. CAHS D is largely disordered, existing in an ensemble of states that resemble a dumbbell composed of two collapsed terminal regions held apart by an extended helical linker^41–43^. We find that the highly helical linker region of CAHS D promotes protection of LDH. Furthermore, the tandem duplication of the linker flanked by endogenous N- and C-termini results in a variant with increased protective capacity relative to wildtype CAHS D. Replacement of the linker region with an exogenous sequence with reduced helicity or scrambling of CAHS D’s endogenous sequence results in reduced helicity and a concomitant decrease in protection. Finer scale disruption of the linker’s helicity via charge identity swapping results in correlated decreases in protection as does the insertion of helix-breaking prolines or the replacement of hydrophobic residues with serine/glycine. In every variant that decreased helicity of CAHS D or its linker region we observed a decrease in protective capacity. Furthermore, we find that the helicity of variants generated by a similar modality (e.g., insertion of prolines) correlates well with protective capacity. However, this correlation decreases when comparing variants generated through different modalities (e.g., insertion of prolines *versus* sequence scrambling), implying that helicity is mechanistically linked to desiccation protection in CAHS D, but is not the only factor governing this phenomenon.

## Results

### CAHS D is a representative CAHS protein

To address the question of whether the helicity of desiccation-related IDPs is necessary for promoting their protective capacity during drying, we chose to study CAHS proteins. This decision was made since these CAHS proteins are known to contain transient helical structure which increases upon desolvation^28^, and they confer protection during drying^29, 44–46^.

One of the best-characterized CAHS proteins is CAHS D. CAHS D has been shown to be required for robust desiccation tolerance in tardigrades, to confer desiccation tolerance to labile enzymes *in vitro*, and to promote anhydrobiosis when exogenously expressed in yeast and bacteria^29^. Biophysical and computational studies have shown that CAHS D occupies a dumbbell-like ensemble of interconverting conformations composed of two relatively compact termini bridged by an internal linker region (Fig. 1A)^41–43^. The terminal regions of CAHS D contain transient helical and beta structure, while the linker region interconverts between random coil and helical structures^41^ (Fig. 1A). The CAHS linker contains both polar and hydrophobic residues, and when in a helical state, this region exists as an amphipathic helix, as quantified by the hydrophobic moment (Fig. 1A).

**Figure 1:**
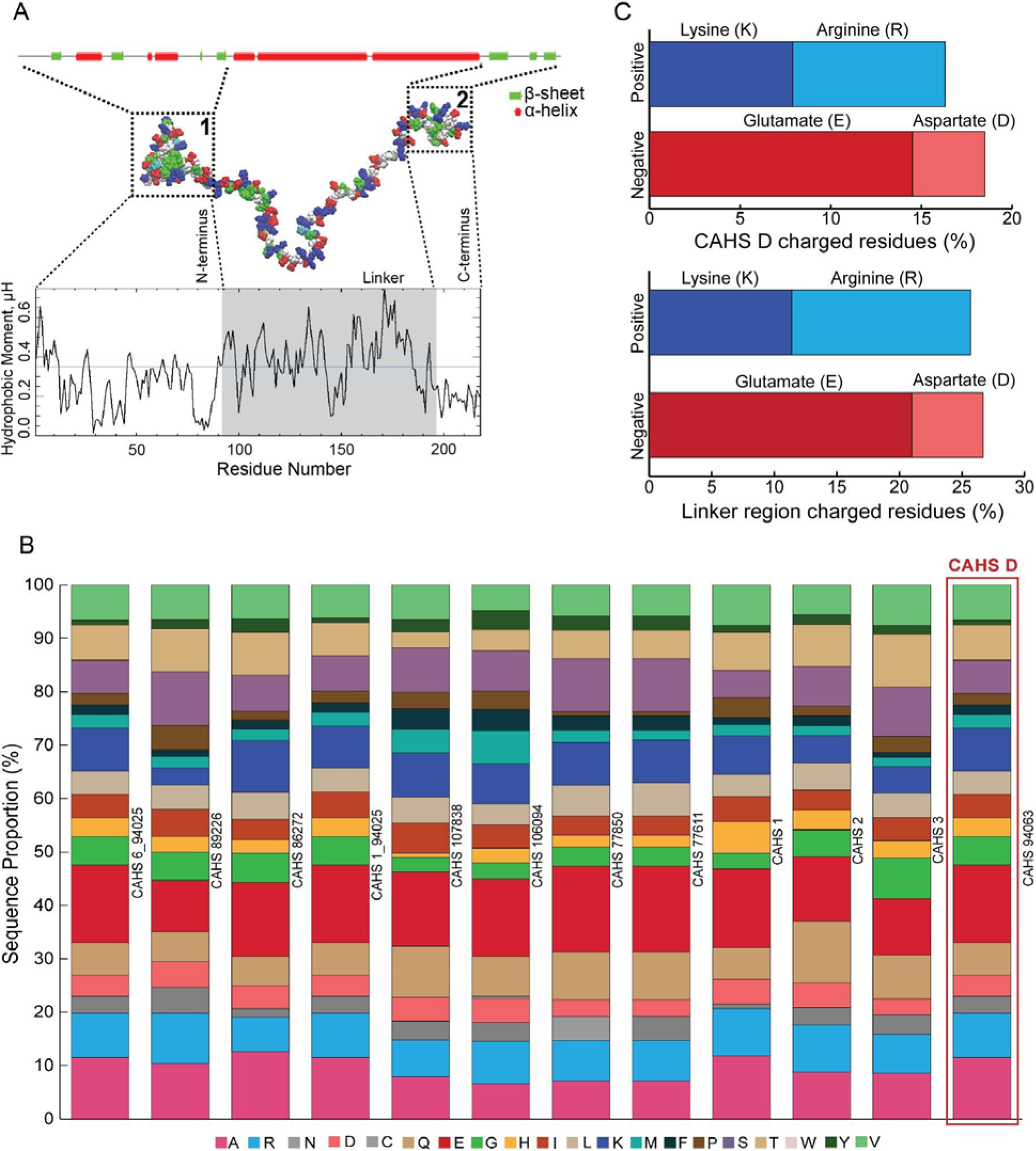
CAHS D, a model CAHS protein. A) Representative conformational ensemble of model CAHS D protein by all atom Monte Carlo simulation^41^. Helix/sheet assignments made using JPRED4^73^. CAHS D hydrophobic moment per amino acid residue calculated by EMBOSS Hydrophobic Moment Calculator^71^. **B)** Amino acid composition of CAHS proteins. **C)** Percentage of charged residues in full length CAHS D (top) and the CAHS D linker region (bottom).

Disordered proteins can be classified by sequence properties, including the fraction of charged residues (FCR), net charge per residue (NCPR), and hydropathy^47^. With this in mind, CAHS D resembles other CAHS proteins in terms of these and other properties, including FCR, NCPR, hydropathy, disorder content, protein length, and overall amino acid composition. CAHS proteins, including CAHS D, have high FCR (Fig. 1B) and a low kappa (_), a measure of charge distribution^41^ (Table 1). Beyond a high FCR, CAHS D is representative of other CAHS proteins in that an approximately equal proportion of positively charged amino acids; lysine (K) and arginine (R) are present, while there is a bias between negatively charged residues; more glutamate (E) as compared to aspartate (D) (Fig. 1B&C). Furthermore, computational analysis (Metapredict and Alphafold2) shows that at the level of ensemble, CAHS D is predicted to resemble other CAHS proteins in the UniProt database, which are predicted to have helical structure in an internal region spanning amino acids ∼91 to ∼195 (Table 1). An exception to this is CAHS 89226, which is larger than other CAHS proteins (414 aa) and whose internal helical linker region is correspondingly offset (∼280-380 aa) (Table 1). Thus, since its sequence and ensemble features mirror those of other CAHS proteins, CAHS D represents a good model for studying the properties and mechanisms underlying CAHS protein function. Furthermore, since CAHS D is protective during desiccation and is known to adopt transient helical structure, especially within its linker region, it is a good model for testing the link between an IDPs helicity and protective capacity during drying.

**Table 1:**
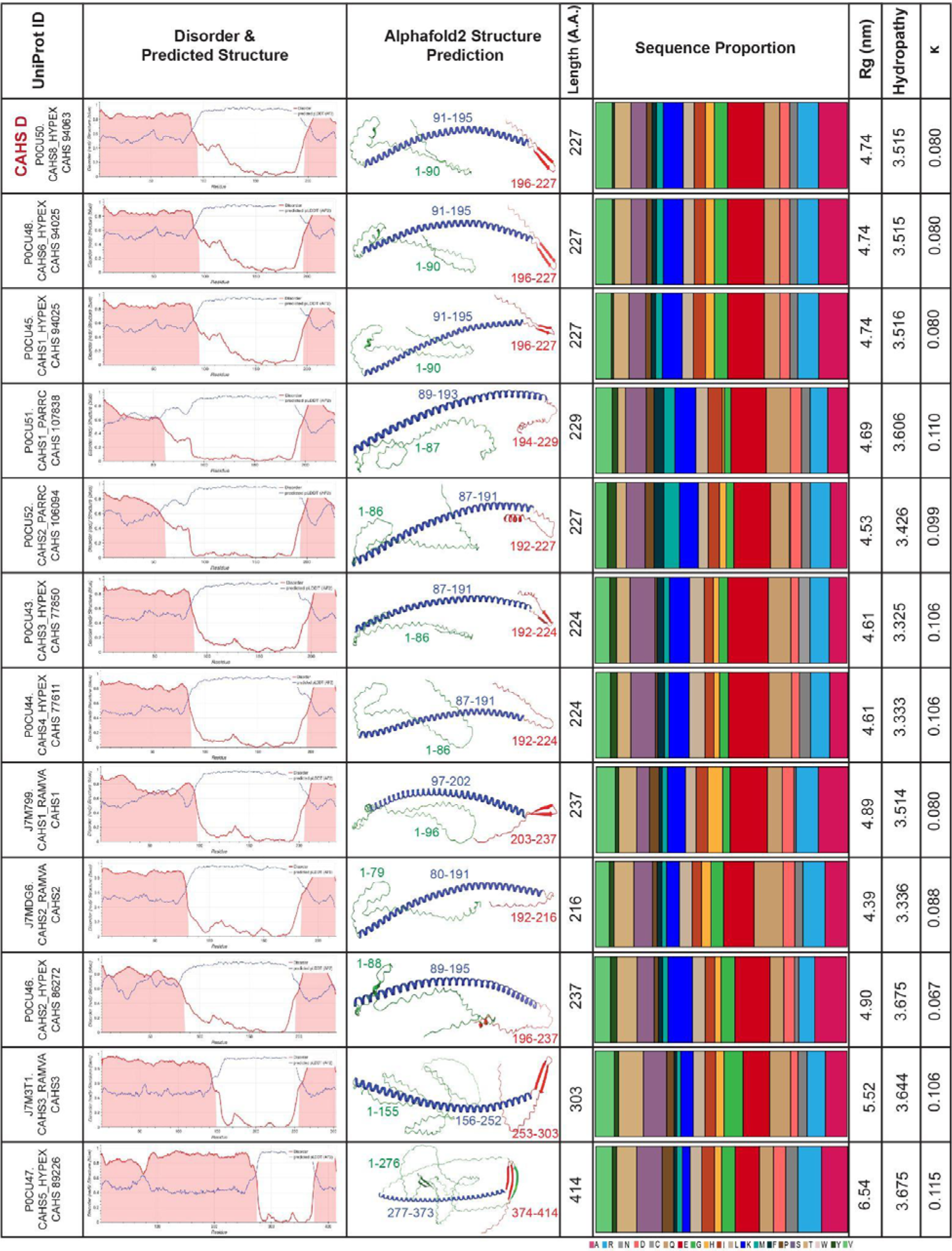
Physical and Structural parameters of CAHS proteins.

### The helical linker region of CAHS D promotes protection of lactate dehydrogenase during drying

To begin to address whether the helicity of CAHS D is important in mediating protection during drying, we tested the ability of the wildtype protein as well as each of its three individual domains (the N-terminus, the helical linker region, and the C-terminus) to stabilize a labile enzyme, lactate dehydrogenase (LDH) during drying (Fig. 2A). We observed that the wildtype protein as well as each of its isolated domains displayed concentration-dependent protection of LDH (Fig. 2B). We next calculated a protective dose 50% (PD50, mM) for the wildtype protein and each of its isolated domains. The PD50 is the concentration at which a protectant preserves 50% of LDH’s enzymatic activity, so a higher PD50 indicates lower protective capacity. We observed that the PD50 of these for protectants varied dramatically (Fig. 2C). Relative to CAHS D, the N- and C-termini show significant decreases in protection (increases in PD50) of ∼8X and ∼7X, respectively, while the protection conferred by the linker region decreases by ∼3X (Fig. 2C). Thus, while no single domain accounts for the entire protective capacity of the wildtype protein, of all three domains of CAHS D the linker region stabilizes LDH the best in isolation.

**Figure 2:**
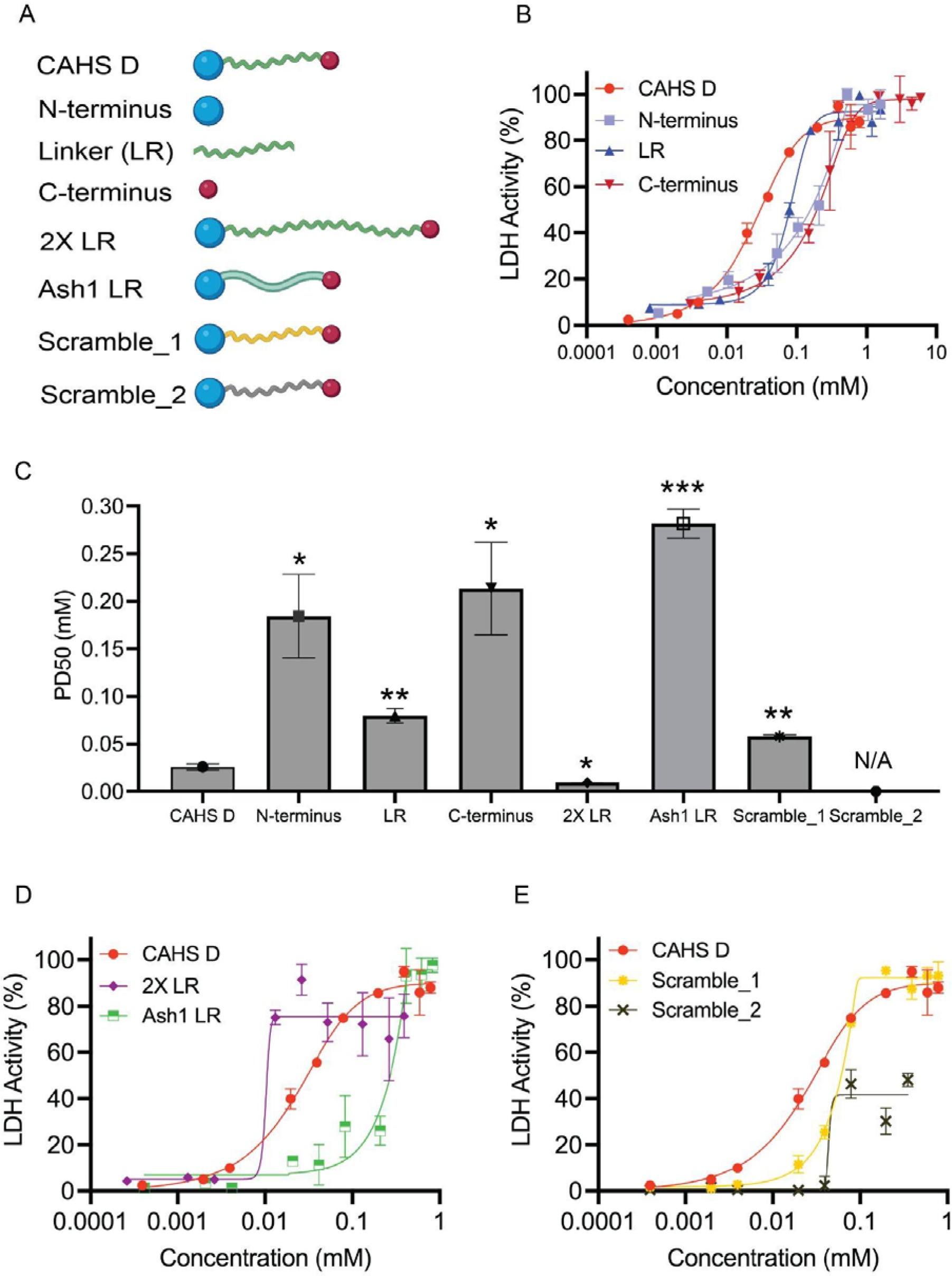
CAHS D Linker Region outperforms other domains in protein unfolding protection. A) Graphical representation of CAHS D domain evaluation variants. **B)** Concentration dependence of LDH protection by CAHS D domains. **C)** 50% protective dose (PD50, mM) for CAHS D variants. **D&E)** Concentration dependence of LDH protection by CAHS D variants. LR = Linker region. n = 3, error bars represent standard deviation (SD).

To further assess the contribution of the helical linker in the context of the full-length protein (e.g., with an N- and C-terminus), we utilized two variants, 2X LR and Ash1 LR (Fig. 2A). 2X LR, is a variant of CAHS D generated by the tandem duplication of the helical linker region. Ash1 LR is a variant generated by replacing the endogenous linker of CAHS D with a duplicated portion of the Ash1 domain of ASH1 (UniProt P34233), a protein involved in transcriptional regulation in *S. cerevisiae* with no connection to desiccation tolerance^48^. This insertion of Ash1 sequence results in a variant of 227 amino acids, the same length as wildtype CAHS D (File S1). In both the 2X LR and Ash1 LR variants, the altered linker regions are flanked by the endogenous N- and C-termini of CAHS D. Consistent with the endogenous helical linker region promoting protection during drying, the 2X LR variant’s PD50 was significantly lower than CAHS D, indicating that doubling the linker results in significantly better protection (Fig. 2C&D). This is in contrast to the Ash1 LR, which provided significantly worse protection compared to wildtype CAHS D (Fig. 2C&D).

Next, we asked whether the composition of the CAHS D linker alone drives protection or whether there is an underlying sequence grammar (e.g., the order of amino acids) that is important for protection. To this end, we utilized two variants of CAHS D that maintain the wildtype sequence composition but change the order and organization of residues within the linker region. These variants, termed here Scamble_1 and Scramble_2 (Fig. 2A; File S1), showed modest concentration-dependent protection (Fig. 2E). Scramble_1 has significantly reduced protection relative to wildtype CAHS D, indicating a partial loss of function (Fig. 2C&E). Scramble_2 failed to reach 50% protection under the concentrations tested, and thus a PD50 could not be calculated, indicating that Scramble_2 has lost even more functionality than Scramble_1 (Fig. 2C&E).

Combined, these experiments identify the helical linker region of CAHS D as the major driver of desiccation protection of the protein. Furthermore, they imply that the sequence composition of CAHS D is insufficient to confer protection during drying.

### Helicity of CAHS D domains and variants correlates with their protective capacity

We wondered what properties of our variants described in Figure 2 correlate with protective capacity during desiccation. To begin to assess this, we asked whether the length of a variant correlates with its protection (Fig. 3A). While there is a strong correlation between the length of variants containing endogenous CAHS D sequence and protective capacity (R^2^ = 0.81; Fig. 3A, blue), we observed that the Ash1 LR variant, which contains exogenous sequence, weakened this correlation (R^2^ = 0.26; Fig. 3A, red). This observation implies that beyond length, there is some property(s) that promote CAHS D-mediate protection.

**Figure 3:**
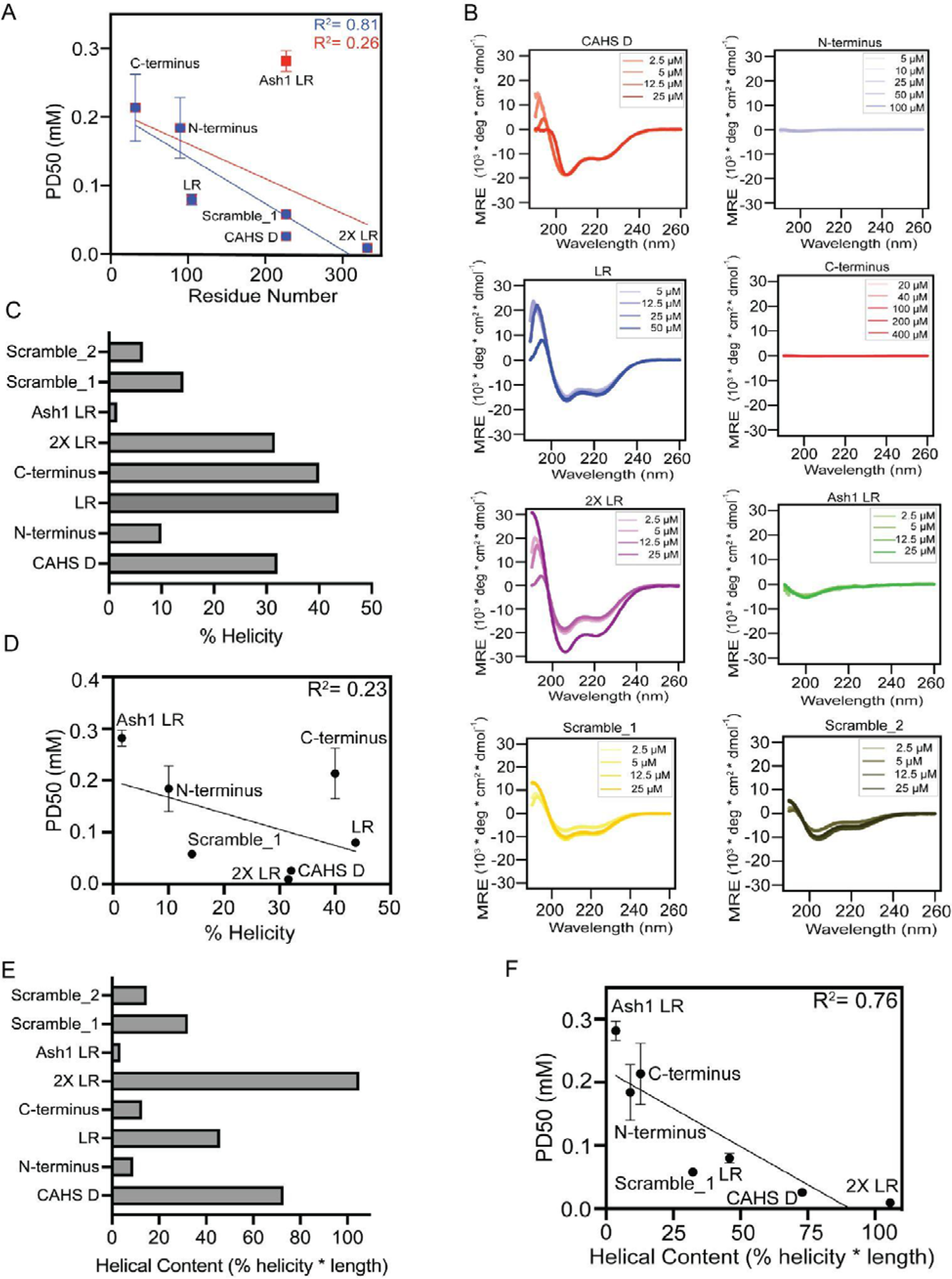
Helicity in the Linker Region is a potential property for protection. A) Amino acid number vs PD50 (mM) correlation plot. **B)** CD spectra for CAHS D domain evaluation variants over a range of concentrations. **C)** Percent helicity deconvoluted from CD spectra from B. **D)** Correlation between percent helicity and PD 50 (mM). **E)** Helical content (% helicity * length) of different CAHS D variants. **F)** Correlation between PD50 (mM) and helical content. All error bars represent Mean±SD.

Since CAHS D is known to be partially helical and helicity has been proposed to be linked to the protective capacity of IDPs during drying^39^, we asked if the helicity of our variants correlates with protection. To test this, we first performed circular dichroism (CD) spectroscopy on each of our variants (Fig. 3B). Spectra were obtained over a 10- fold concentration range to test for structural change that may occur due to self association. However, no significant structural change is seen for any of the variants in the concentration range tested (Fig. 3B). Raw CD spectra were deconvolved allowing us to derive an estimate of the percent helicity of each variant (3C). Correlating percent helicity to protection did not result in a compelling trend (R^2^ = 0.23; Fig. 3D).

Seeing that percent helicity and protection do not correlate well might imply that helicity is unimportant for protection. However, we reasoned that protection is likely a function of both a protein’s length and its percent helicity. For example, if two proteins are both 10% helical but one is 10 amino acids and the other 1000, the 1000 amino acid protein will on average have much more helical content. If helicity is important for protection during drying then both percent helicity and the length of a protein likely need to be taken into account.

To this end, we calculated the helical content (percent helicity x length) for each variant. These calculations revealed a decrease in the helical content in Ash1 LR as compared to CAHS D and linker region (Fig 3E). Consistent with the linker region containing a significant degree of helicity, our 2X LR variant contained two times the amount of helical content as the linker region alone (Fig. 3E). Furthermore, terminal regions of CAHS D in isolation \display lower levels of helical content as compared to the wildtype protein or the linker region (Fig. 3E). Finally, the helicity content of Scramble_1 and 2 was decreased relative to wildtype CAHS D (Fig. 3E). Relating the protective capacity (PD50) of each of these CAHS D variants with their helical content resulted in a robust correlation (R^2^ = 0.76) (Fig. 3F).

Taken together, these results suggest that the helicity of CAHS D may help mediate protective capacity during drying.

### Helicity of the CAHS D linker is necessary for robustly protecting lactate dehydrogenase during desiccation

We sought to further probe the link between helicity of the linker region of CAHS D and its protective capacity during drying by generating mutants specifically designed to reduce helical content while maintaining as much endogenous sequence composition and grammar as possible. These mutants consist of either the full-length CAHS protein or linker region in isolation with helix breaking proline residues replacing endogenous residues at every 5-8 residues within the linker region (File S1). These variants we term full-length linker region proline (FL_LR_P) and linker region proline (LR_P), respectively. In these variants, the replacement of endogenous residues with proline maintains total amino acid number, but modestly increases isoelectric point and molecular weight of the resulting protein (Table 2).

**Table 2:** Physicochemical features of CAHS D variants used in the study.

| **Variants** | **AA Residue Number** | **Isoelectric Point (p<sup>I</sup>)** | **MW (Da)** | **Helical Content (% helicity \* length)** |
| --- | --- | --- | --- | --- |
| CAHS D | 227 | 5.98 | 25485.32 | 72.867 |
| N-terminus | 90 | 6.02 | 9547.62 | 9 |
| LR | 105 | 6.07 | 12620.11 | 45.885 |
| C-terminus | 32 | 5.45 | 3353.62 | 12.8 |
| 2X LR | 332 | 6 | 38160.47 | 105 |
| Ash1 LR | 227 | 10.84 | 24119.87 | 3.632 |
| Scramble_1 | 227 | 5.98 | 25485.32 | 32.234 |
| Scramble_2 | 227 | 5.98 | 25485.32 | 14.75 |
| FL_LR_P | 227 | 6.12 | 25652.75 | 6.356 |
| LR_P | 105 | 6.91 | 12787.54 | 2.73 |
| FL_LR_NH | 227 | 5.98 | 24845.60 | 2.724 |
| LR_NH | 105 | 6.06 | 11980.39 | 1.05 |
| LR_KtoR | 105 | 6.07 | 12956.27 | 41.475 |
| LR_RtoK | 105 | 6.07 | 12199.91 | 19.11 |
| LR_DtoE | 105 | 6.10 | 12690.25 | 31.71 |
| LR_EtoD | 105 | 5.85 | 12311.52 | 7.98 |

We began characterizing FL_LR_P and LR_P variants by assessing their helical content using CD spectroscopy (Fig. 4A&B). Consistent with previous reports that prolines are potent disruptors of secondary structure^49, 50^, we observed a decrease in helical content to negligible levels in both our FL_LR_P and LR_P variants (Fig. 4B). Here, as before, no change in CD spectra was observed over the range of concentrations tested (Fig. 4B).

**Figure 4:**
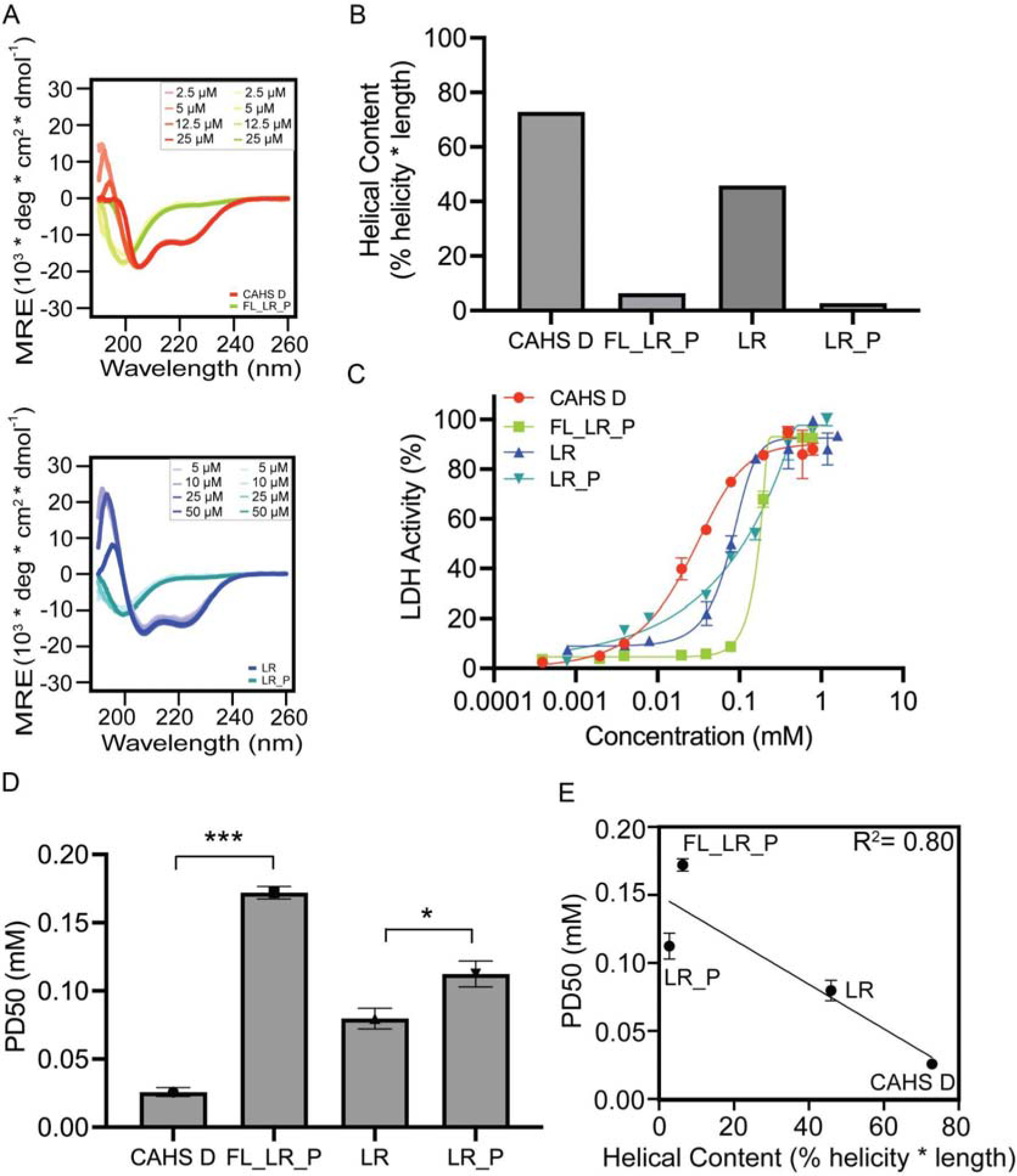
Helix disrupting sequence variations disrupt protection. A) CD spectra of full length (top) and linker (bottom) proline insertion variants, as well as wildtype proteins. **B)** Helical content deconvoluted and normalized from CD spectroscopy data. **C)** Concentration dependence of proline insertion variants in protecting LDH during drying. n = 3, error bars represent standard deviation (SD). **D)** PD50 value calculated from fitted sigmoidal curve from Fig. 4C **E)** Correlation between PD50 (mM) and helical content (% helicity * length). All error bars represent Mean±SD. Welch’s t-test, ns = not significant, * ≤ 0.5, ** ≤ 0.1, *** ≤ 0.05

Next, we performed LDH desiccation assays to assess how direct disruption of helicity within the context of wildtype CAHS D, or the linker region in isolation, affects the protective capacity of these IDPs. In both cases, we observed significant decreases in the protective capacity of FL_LR_P and LR_P relative to both wildtype CAHS D and isolated linker region, represented by a significant increase in the PD50 value for each variant (Fig. 4C&D). Furthermore, the correlation of the helical content of wildtype CAHS D, its linker region, and the proline variants with protective capacity shows a robust correlation with an R^2^ value of 0.80 (Fig. 4E).

Taken together, these results suggest that the helicity of CAHS D, and in particular the linker region of CAHS D, is necessary for robust protection during drying.

### Sequence identity of charged residues in the CAHS D linker promotes helical structure and protection during drying

After observing that direct disruption of helicity within the linker region of CAHS D results in a loss of robust protective capacity during drying, we were curious what sequence features within the linker promote helicity. To this end, we examined the primary sequence of CAHS D and its linker more closely.

Analysis of the linker region sequence using CIDER^51^ reveals a high fraction of charged residues (FCR= 0.524) and characterizes this domain as a strong polyampholyte (sequence containing positively and negatively charged amino acids)^47^. The amino acid sequences of polyampholyte helices have been found to have a high amount of glutamate (E) and both positively charged lysine (K) and arginine (R) residues in a regular, repeating pattern of oppositely charged residues^52^. Consistent with this, out of the high fraction of charged residues in the linker region of CAHS D, there is an approximately equal amount of positively charged K and R (12:15) residues whereas for negatively charged amino acids, there is a bias for E relative to D (22:6) residues (Fig. 1B&C). Thus, the ratio and identities of charged residues within the linker region of CAHS D mirrors that of traditional polyampholyte helices.

To test the contribution of charge identity to the helical nature of CAHS D’s linker region as well as to desiccation protection, we generated four charge identity variants. These variants, called LR_KtoR, LR_RtoK, LR_EtoD, and LR_DtoE, were generated by replacing all the Ks for Rs (and vice versa) or all the Es for Ds (and vice versa) within the isolated linker region. These mutants maintain the length and charge of the linker region and allow us to further test the relationship between sequence features, helicity, and protective capacity of the CAHS D linker region during drying.

To begin, we tested the effect of these charge identity variants on helicity using CD spectroscopy. CD spectroscopy revealed that changing the identity of charged residues, while still maintaining overall charge influences the helicity of the linker region (Fig. 5A&B). Specifically, disruption of the highly biased presence of E, by replacement with D, showed a decrease in helicity content in LR_EtoD as compared to the wildtype linker region. The converse was also true, as replacing the relatively few D residues in the linker region with E residues (LR_DtoE) also reduced the helical content. However, LR_DtoE maintained more helicity than LR_EtoD, which is consistent with a recent study suggesting that glutamate helps maintain helicity, while aspartate helps maintain extended structures in IDPs^53^. This trend partially carried over to positively charged residues as well, with our LR_RtoK variant showing reduced helical content, while the helical content of the LR_KtoR variant remained more or less consistent with that of the wildtype linker region.

**Figure 5:**
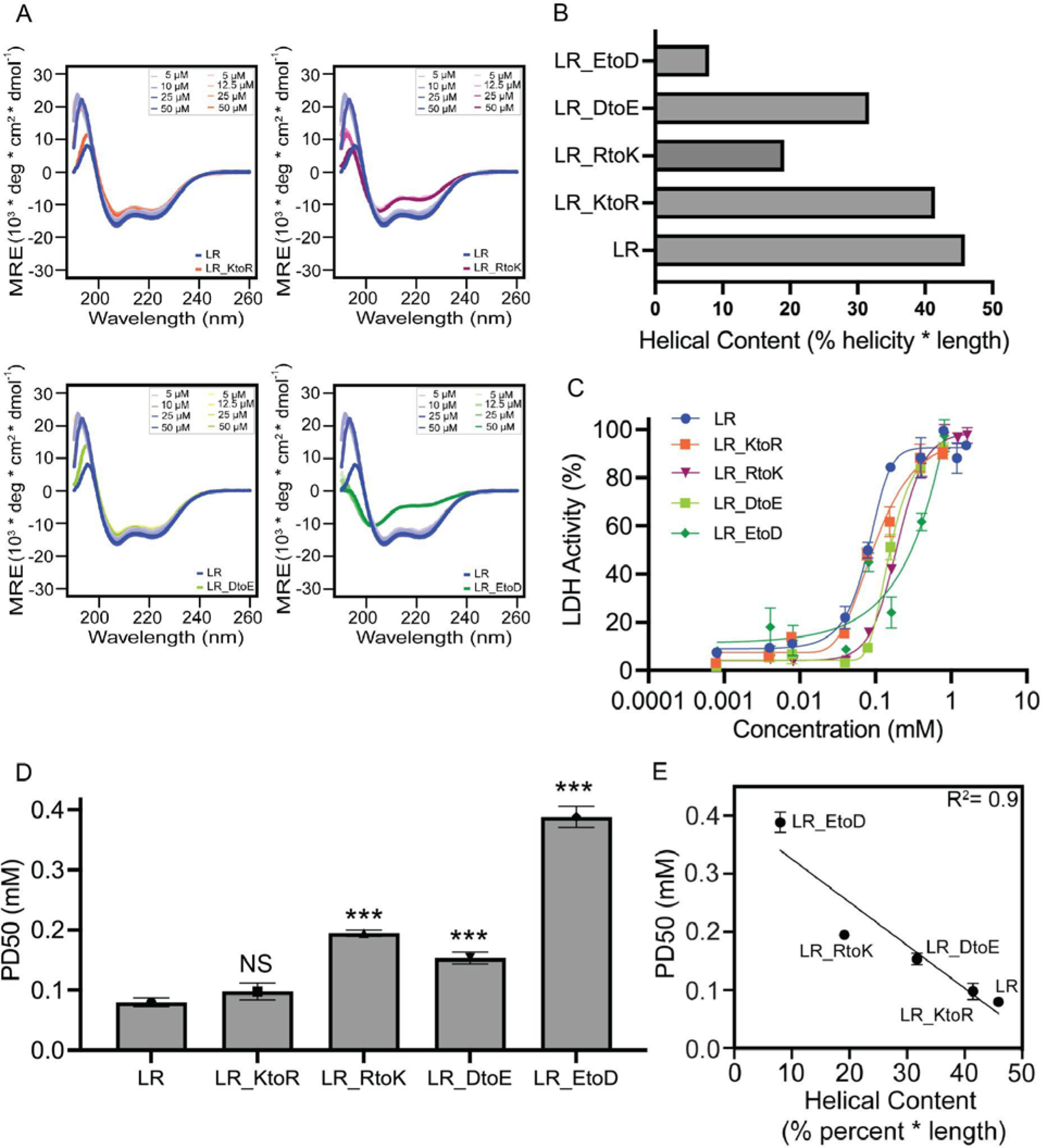
Charge identity defines fine tuning of helix formation in CAHS D Linker Region. A) CD spectra of CAHS D Linker Region charge identity variants. **B)** Helical content deconvoluted and normalized from CD spectroscopy data. **C)** Sigmoidal curve representing LDH unfolding protection. Protection is plotted as the mean percentage of LDH enzyme activity against a range of concentrations in mM. n = 3, error bars represent standard deviation (SD). **D)** PD50 (mM) calculated from the fitted sigmoidal curve from Fig. 5C. **E)** Correlation between PD50 and helical content. All error bars represent Mean±SD. Welch’s t-test, ns = not significant, * ≤ 0.5, ** ≤ 0.1, *** ≤ 0.05

Next we tested the protective capacity of these four charge identity variants using the LDH assay. We observed significant decreases in protective capacity for all charge identity variants except LR_KtoR (Fig. 5C&D). Correlative analysis comparing the helical fraction and PD50 of these variants revealed a strong correlation (R^2^= 0.9) between these properties (Fig. 5E).

These results demonstrate that charge identity contributes to helicity of the CAHS D linker region. Furthermore, these results provide further evidence that helicity is linked to the protective capacity of the CAHS D linker during drying.

### Removal of hydrophobic residues does not rescue loss-of-helicity variants

Our results thus far implicate helicity as an important feature in determining the protective capabilities of CAHS. One possible interpretation is that transient helicity ensures CAHS linker residues are exposed along helical faces and therefore primed for inter-molecular interactions with clients such as LDH. Transient intra-molecular interactions driven by hydrophobic residues can lead to IDR compaction, which minimizes the potential for intermolecular interactions^54, 55^. If this is the case, loss of helicity may lead to aberrant intra-molecular interactions driven by hydrophobic residues, which in turn could impede inter-molecular interactions and the protective capabilities of the linker region.

To test this hypothesis, we designed variants to simultaneously disrupt helicity and remove hydrophobic residues by replacing hydrophobic residues with either serine or the helix-breaking residue glycine. Variants of both the linker in isolation (LR_NH) and the linker in its endogenous context (FL_LR_NH) were designed. As predicted, these designs led to a dramatic decrease in helical content (Fig. 6A&B; File S1). Importantly, similar to our previous observations, we also measured a significant decrease in the protective capacity of FL_LR_NH compared to wildtype CAHS D as well as LR_NH compared to the linker region in isolation (Fig. 6C-E). These results further support a model in which helicity is critical for CAHS protective capabilities and suggest that aberrant intra-molecular interactions driven by hydrophobic residues do not underlie the reduced protection observed in low-helicity sequences.

**Figure 6:**
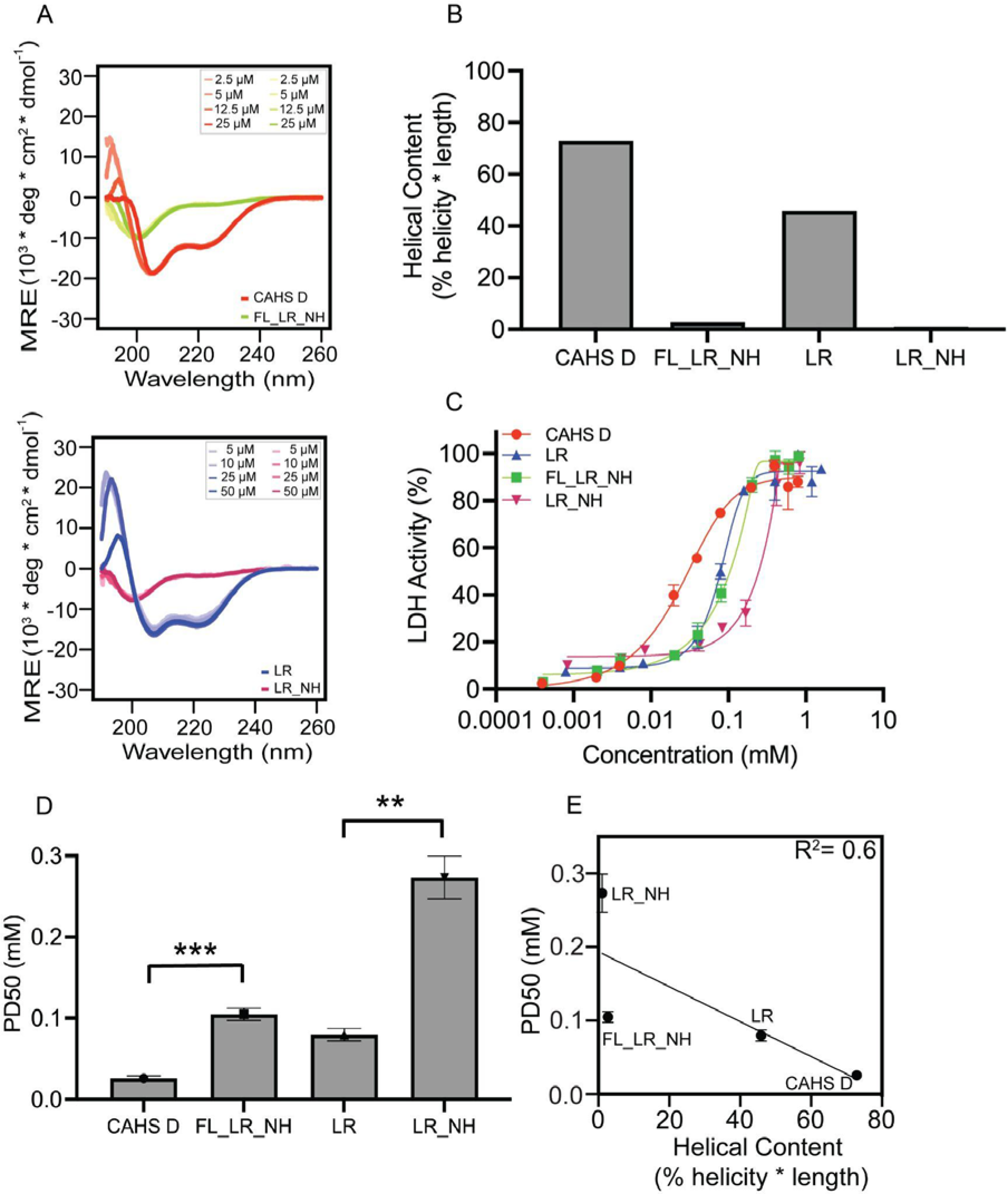
Hydrophobic residues help in helix formation and desiccation protection. A) CD spectra for no hydrophobic variants, as well as wildtype proteins. **B)** Helical content deconvoluted and normalized from CD spectroscopy data. **C)** Sigmoidal curve representing LDH unfolding protection. Protection is plotted as the mean percentage of LDH enzyme activity against a range of concentrations in mM. n = 3, error bars represent standard deviation (SD). **D)** PD50 value calculated from the fitted sigmoidal curve. **E)** All error bars represent Mean±SD. T-test, ns = not significant, * ≤ 0.5, ** ≤ 0.1, *** ≤ 0.05

## Discussion

An emerging paradigm in the anhydrobiosis field is the prevalent enrichment of IDPs that help to mediate protection during drying^19, 26, 33^. Transient helicity, and the acquisition of increased helical content during desolvation, has been reported in many desiccation-related IDPs spanning diverse families^20, 28, 37, 56^. Helical structure has often been speculated to be essential for desiccation-related IDP function during drying, but direct, empirical evidence demonstrating this linkage is lacking. Here we show that helical structure found in the linker regions of CAHS D, a desiccation-related IDP required for tardigrades to survive drying, is necessary to robustly protect the labile enzyme LDH during drying.

### The helical linker region is the protective domain of CAHS D

CAHS D has previously been shown to be essential for robust anhydrobiosis in the tardigrade *H. exemplaris*, to increase desiccation tolerance in heterologous systems, and to be sufficient to protect labile enzymes *in vitro*^29^. Furthermore, CAHS D occupies a dumbbell-like ensemble of interconverting conformations, composed of two collapsed termini held apart by an extended linker region. However, previous studies have not examined which of these three domains contribute to the protective capacity of CAHS D. Nor have they examined what sequence features, including helicity, are important for CAHS D-mediated desiccation tolerance.

In our current study, we show that while wildtype CAHS D more efficiently protects LDH than any of its three domains in isolation, the linker region has significantly more protective capacity than either terminal region. Furthermore, replacement of the linker region with an exogenous sequence or with scrambled sequences that conserve endogenous composition by altering grammar results in large decreases in protection. Conversely, a tandem duplication of the linker region results in a variant with increased protective capacity relative to the wildtype protein. From this we infer that the domain which most prominently drives CAHS D mediated protection is the internal linker.

### Molecular grammar underlying CAHS D linker helicity

Replacing the endogenous linker of CAHS D with a novel sequence or altering the sequence of the linker either by conserving composition, but scrambling the order of amino acids, swapping charged amino acid identities, or by replacement of hydrophobic residues with helix-breaking glycines or serines results in significant decreases in protective capacity. These experiments imply that not only is the composition of the linker important for protection, but so too is the molecular grammar of this domain.

Coupled with biophysical assessment of secondary structure, an overarching conclusion from these sequence variants and functional studies is that sequence composition and grammar promoting helicity also promotes protection.

The observed molecular grammar underlying CAHS D linker helicity is in line with previous reports. For example, *in silico* analysis of sequences containing glutamate (E) are predicted to have greater helicity than those containing aspartate (D) (*e.g.,* (E_4_K_4_)_2_ *versus* (D_4_K_4_)_2_), while sequences containing arginine have a higher predicted helicity than those containing lysine (*e.g.,* (D_4_R_4_)_2_ versus (D_4_K_4_)_2_)^52^. Within the linker of CAHS D there is a clear bias for negatively charged E relative to D (22:6), which should promote helicity. Consistent with this, converting all E residues to Ds within the linker (LR_EtoD) results in a dramatic loss of helicity (and protection) relative to converting D to E (LR_DtoE). Whereas, converting all K residues to R (LR_KtoR) has little effect on helical content, while the converse (LR_RtoK) dramatically decreases linker helicity.

### Helicity and desiccation protection

Our study demonstrates that the protective capacity of CAHS D is linked to helicity. Previous work has shown that during desiccation or chemical desolvation CAHS proteins increase in helical structure^28^, suggesting that drying may induce functionality. This is in line with previous reports that other desiccation-related IDPs, for example some LEA proteins increase in helical content upon desiccation^20, 56, 57^.

Our observation that perturbing helicity of the CAHS D linker *via* the insertion of prolines, swapping of charged amino acids, removal of hydrophobic residues or through sequence scrambling results in decreased protective capacity during drying provides insights into the necessity of these structural and biophysical properties not only for CAHS proteins, but also for other stress-related IDPs such as LEA proteins.

### Helicity drives, but is not the only property contributing to, desiccation tolerance

In every perturbation we performed that decreased the helicity of CAHS D or the CAHS D linker we observed a decrease in protective capacity. Furthermore, correlating the helical content of variants generated through a single modality (e.g., charge identity swapping) with protection resulted in robust correlations. However, combining all these variants onto a single plot dramatically decreases trend, as does grouping all full-length and linker region variants individually (Fig. 7A-C). Thus, while helicity of CAHS D may be linked to its protective function, helicity alone is not sufficient to accurately predict absolute protective capacity. This prompts the obvious question of what property(s) of a helix make it protective or non-protective during desiccation.

**Figure 7:**
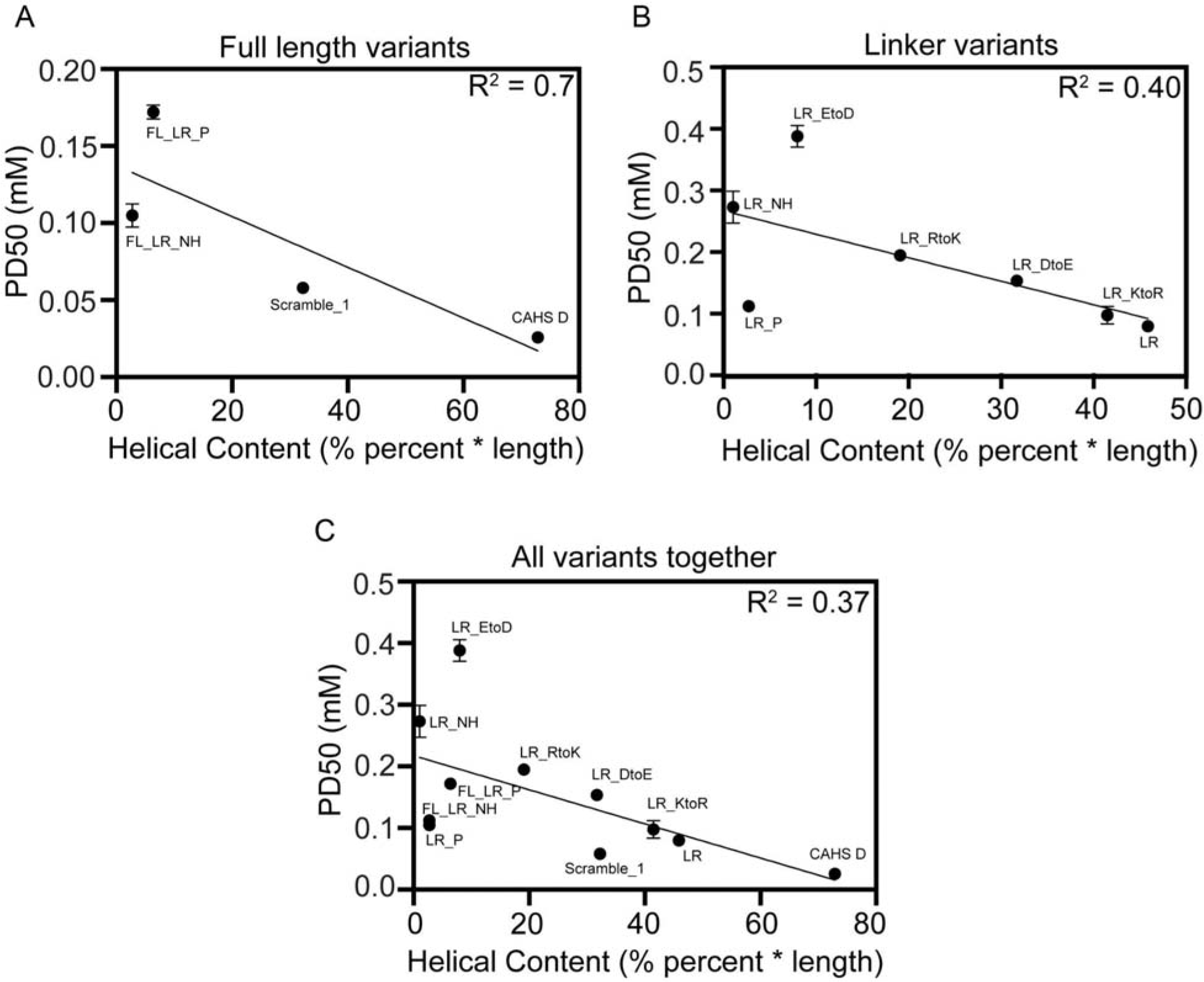
Correlation of protective dose 50 and helical content for all variants used in this study. A) Correlation of PD50 and helical content for full length variants. **B)** Correlation of PD50 and helical content for linker variants. **C)** Correlation of PD50 and helical content for all variants. All error bars represent Mean±SD.

One commonality shared between the helices formed by the CAHS D linker and other desiccation related IDPs, such as LEA proteins, is their amphipathic nature. Amphipathicity of LEA proteins has previously been proposed to allow for hydrophobic interactions between LEA proteins and client molecules (e.g., membrane headgroups) and could play a role in self-assembly of proteins into oligomers^58, 59^. While no evidence exists that CAHS D, or its linker region, interact with membranes, several studies have demonstrated that CAHS D, as well as other CAHS proteins, oligomerize^41, 43, 46, 60^.

Furthermore, the helicity and amphipathic nature of the linker region of CAHS D has been shown to be essential for oligomerization^41^.

Perhaps an overarching theme for desiccation-related IDPs that rely on helicity to promote protection is that their oligomerized state relies on the adoption of amphipathic helices, and that self-assembly into an oligomerized state promotes protection during desiccation. But, how might oligomerization promote protection? LEA proteins have been shown to prevent aggregation of proteins, both *in vitro* and *in vivo*^20, 61, 62^. Along these lines, LEA proteins are thought to serve as molecular shields, molecules that sterically prevent the association of aggregation protein prone proteins during drying^61–63^. An oligomerized state might promote molecular shielding by increasing the radius of gyration of LEAs, allowing them to take up more room and work more efficiently at preventing aggregation.

Alternatively, oligomerization might result in altered surface chemistries. Amphipathic helices often associate *via* hydrophobic interactions^64^. Dimers or higher order oligomers formed from amphipathic helices through hydrophobic interactions could result in complexes with increased surface hydrophilicity, which could in turn promote association and interaction with residual water, creating local areas of hydration, or alternatively might serve to prevent the unfolding of proteins by creating a hydrophilic environment.

### Beyond helicity and secondary structure

While many desiccation-related IDPs have been shown to possess transient helicity, which increases upon drying, this is by no means an absolute rule. For example, SAHS proteins from tardigrades contain more stable beta-structure^65^, as do some LEA proteins^66^. Even CAHS proteins are known to contain transient beta-structure, albeit localized to their terminal regions. How transient secondary structure beyond helices influences or promotes the protective capacity of IDPs during drying is unknown.

As touched upon above, many desiccation-related IDPs are known oligomerize. The adoption of quaternary structure, either through helical interactions or other means, is likely to also play a role in promoting protection during drying. CAHS D provides a compelling example of the potential role of quaternary structure in IDP-mediated anhydrobiosis. CAHS D is known to form higher order oligomers in a concentration dependent fashion, and mutagenesis disrupting this oligomerization has been seen to augment the protective capacity and mechanism of this protein^41^.

Thus, while our study shows empirically that helicity of CAHS D, a desiccation-related IDP, is linked to its protective capacity, it also demonstrates that other properties, likely play a role in CAHS D-mediated desiccation tolerance.

## Materials and methods

### Sequence analysis

CAHS proteins’ sequences were obtained from the UniProt database (https://www.uniprot.org/). Protein parameters were computed using Expasy ProtParam tool^67^. Intrinsic disorder and predicted structure was predicted using metapredict online (v2.3)^68^. Parameters related to disordered protein sequence such as kappa (_) value, hydropathy, and diagram of states were calculated using CIDER^51^. Protein structure was predicted by Alphafold2^69^ and visualized using ChimeraX^70^. Hydrophobic moment of the CAHS D sequence was calculated and plotted using the EMBOSS hmoment calculator^71^.

(https://www.bioinformatics.nl/cgi-bin/emboss/hmoment)

### Sequence design tool: Charge identity mutants, hydrophobic residue mutants

Synthetic disordered proteins were designed using GOOSE, a tool for the rational design of synthetic disordered proteins (https://github.com/idptools/goose). In all cases, disorder was revaluated using metapredict^68^, and designs were generated using GOOSE’ variant design feature.

### CAHS D variants’ construct design, cloning, and -80C freezer stock preparation

*E.coli* codon optimized gBlocks encoding CAHS D and its mutant proteins used in this study were synthesized by (Integrated DNA Technologies) and cloned into pET-28 b (+) vector (Addgene) for bacterial expression. Plasmids containing CAHS D or its variants were transformed initially into DH5_ *E.coli,* plated onto Kanamycin (Kan) containing LB agar plates. Selected colonies were grown overnight in the LB broth followed by plasmid extraction for the sequence confirmation (Eton Bioscience). Sequence verified plasmids were then transformed into BL21 (DE3) *E.coli* expression strains. BL21 (DE3) transformed colonies from LB+Kan plates were verified by a small scale protein expression test, and successful colonies were used to make -80 _ freezer stocks.

### Expression

10 ml of saturated overnight (16 hr) culture of respective CAHS D variant was used to inoculate 1 L of LB broth (10 g/L peptone, 5 g/L yeast, 5 g/L NaCl) supplemented with Kanamycin to a final concentration of 50 μg/mL. The cultures were shaken (180 rpm) at 37 _ until the optical density at 600 nm was 0.6 (not higher than 0.8). Once the cultures reached the expected OD, a final concentration of 1 mM IPTG was used to induce the target protein expression. After 4 hr, cells were collected as pellets by centrifuging at 4000 rpm for 30 min at 4 _. The cell pellets from 1 L culture were resuspended into 5 ml of 20 mM Tris (pH 7.5) and mixed with 30 μl of 1X protease inhibitor (Sigma-Aldrich).

### Protein Purification

Resuspended cell pellets were heat shocked at 95 _ for 15 min and spun down at 10,500 rpm for 30 min at 10 _ followed by a 0.22 µM syringe filtration (EZFlow Syringe Filter, Cat. 388-3416-OEM). The heat-soluble supernatant was mixed with 2X volume of 8 M urea-containing 50 mM sodium acetate buffer (pH 4.0). The protein sample in urea was further purified on Cytiva AKTA Pure chromatography system (UNICORN 7 Workstation pure-BP-exp) using a cation exchange column (Cytiva HiPrep^TM^, SP HP 16/10, catalog #29018183). The proteins were eluted using gradients of 1 M NaCl. Different CAHS D variants will elute at different salt concentrations. SDS-PAGE was used to identify fractions containing pure CAHS D or its mutant variants. Confirmed fractions were pooled, transferred to a dialysis tubing of 3.5 kDa membrane pore size (spectrumlabs.com) and dialyzed against 20 mM Phosphate buffer (pH 7) for at least 4hr followed by 6 rounds of 4 hr dialysis against Milli Q H_2_O. Dialysate containing purified CAHS D variant was flash-frozen and lyophilized for 48 hr before storage at -20_.

### Lactate dehydrogenase assay

LDH activity assay was based on the procedure outlined by Goyal et al. 2005^61^. Lyophilized CAHS D and its mutants’ were resuspended into 25 mM Tris-HCl (pH 7) and concentrations were quantified using Qubit^TM^ Protein Assay Kit (Catalog number Q33211). Rabbit Muscle L-LDH (Sigma-Aldrich, REF 10127230001) was mixed to a range of sample protein concentrations having a final concentration of 0.1 g/L. The range of sample concentration for CAHS D, FL_LR_P, and FL_LR_NH used in this assay is (0.01-20) g/L, whereas for CAHS D w/ Ash1, LR and all its variants it is (10- 0.01) g/L. The final volume of 0.1 g/L LDH mixed with (20/10- 0.01) g/L test variants is 100 μL. Half of the sample was stored at 4 °C, and the other half was desiccated using speedvac for 16 hr without heating (SAVANT Speed Vac Concentrator). Desiccated samples were rehydrated with 250 μL molecular grade H_2_O, whereas unstressed control samples stored in 4 °C were diluted with 200 μL of molecular grade H_2_O, and all samples were kept on ice until the enzymatic activity was measured. To check the LDH activity, 10 μL of the sample was added to a mixture containing 980 μL of pyruvate phosphate buffer (100 mM sodium phosphate, 2 mM sodium pyruvate; pH 6.0) and 100 µM NADH (Sigma-Aldrich NADH; disodium Salt, grade II) and the enzyme kinetics was measured. LDH enzyme activity in the conversion of pyruvate to lactate is calculated by measuring the loss of NADH at UV absorbance of 340 nm using nanodrop (Thermo Scientific NanoDrop One) at each two seconds for a kinetic loop of 1 min. The activity was determined as a ratio of NADH loss in stressed samples compared to the controls. LDH activity was measured in triplicates for each sample concentration for all different CAHS D variants.

### Circular dichroism spectroscopy

Lyophilised CAHS D and its variants were resuspended in tris buffer pH 7 to a concentration of 25 µM. Protein concentrations were quantified using Qubit™ Protein Assay. The suspension was then diluted with a 25 mM Tris buffer, pH 7 to concentrations ranging from (2.5-25) µM for measurement. Aliquots of each concentration were measured in 1 mm and 0.05 mm quartz cuvettes in a circular dichroism spectrometer (Jasco, J-1500 model). Each measurement was performed in three replicates.

### Deconvolution of circular dichroism spectra and normalization of helical content

Beta Structure Selection method (BeStSel) was used to calculate the detailed structure information from the CD spectra^72^. The deconvoluted percent of _-helix was normalized to the amino acid residue number of different CAHS D variants to calculate the helical content (% helicity * length).

### Data visualization and Statistical Analysis

CD spectroscopy data was plotted using R-Studio v1.3.1073. Sigmoidal curve plotting, analysis of 50% protective dose value followed by data visualization, and regression analysis were done using GraphPad Prism v9.5.1. Statistical difference of PD50 value between variants was analyzed using Welch’s t-test.

## Acknowledgements

We thank members of the Water and Life Interface Institute (WALII), supported by NSF DBI grant # 2213983, for helpful discussions, as well as support from the NSF via the IntBio research program under awards 2128069 to TCB, 2128067 to SS, and 2128068 to ASH. Fellowship to SB funded by Wyoming NASA EPSCoR, NASA Grant #80NSSC19M0061 supported this work. This work was supported by the USDA National Institute of Food and Agriculture, Hatch project #1012152. Anja Thalhammer is thanked for helpful discussions and insights.

---

**File S1: Sequences of all proteins used in this study File S2: Scripts used in this study**

**File S3: Data used in this study**

## References

1. Armstrong, L. E. & Johnson, E. C. Water Intake, Water Balance, and the Elusive Daily Water Requirement. Nutrients 10, (2018).

2. Tunnacliffe, A. & Lapinski, J. Resurrecting Van Leeuwenhoek’s rotifers: a reappraisal of the role of disaccharides in anhydrobiosis. Philos. Trans. R. Soc. Lond. B Biol. Sci. 358, 1755–1771 (2003).

3. Crowe, J. H. & Clegg, J. S. Anhydrobiosis. (1973)

4. Crowe, J. H., Hoekstra, F. A. & Crowe, L. M. Anhydrobiosis. Annu. Rev. Physiol. 54, 579–599 (1992).

4. Sallon, S. et al. Germination, genetics, and growth of an ancient date seed. Science 320, 1464 (2008).

5. Crowe, J. H., Crowe, L. M. & Chapman, D. Preservation of Membranes in Anhydrobiotic Organisms: The Role of Trehalose. Science (1984) doi:10.1126/science.223.4637.701.

6. Trehalose Renders the Dauer Larva of Caenorhabditis elegans Resistant to Extreme Desiccation. Curr. Biol. 21, 1331–1336 (2011).

7. Tapia, H. & Koshland, D. E. Trehalose is a versatile and long-lived chaperone for desiccation tolerance. Curr. Biol. 24, 2758–2766 (2014).

8. Nguyen, K., Kc, S., Gonzalez, T., Tapia, H. & Boothby, T. C. Trehalose and tardigrade CAHS proteins work synergistically to promote desiccation tolerance. Commun Biol 5, 1046 (2022).

9. Hara, Y., Shibahara, R., Kondo, K., Abe, W. & Kunieda, T. Parallel evolution of trehalose production machinery in anhydrobiotic animals via recurrent gene loss and horizontal transfer. Open Biol. 11, 200413 (2021).

10. Cesari, M., Altiero, T. & Rebecchi, L. Identification of the trehalose-6-phosphate synthase (tps) gene in desiccation tolerant and intolerant tardigrades. *Ital*. J. Zool. 79, 530–540 (2012).

11. Ingemar Jönsson & Persson. Trehalose in three species of desiccation tolerant tardigrades. Open Zool. J.

12. Hengherr, S., Heyer, A. G., Köhler, H.-R. & Schill, R. O. Trehalose and anhydrobiosis in tardigrades--evidence for divergence in responses to dehydration. FEBS J. 275, 281–288 (2008).

13. Westh, P. & Ramløv, H. Trehalose accumulation in the tardigrade Adorybiotus coronifer during anhydrobiosis. J. Exp. Zool. (1991).

14. Lapinski, J. & Tunnacliffe, A. Anhydrobiosis without trehalose in bdelloid rotifers. FEBS Letters vol. 553 387–390 Preprint at https://doi.org/10.1016/s0014-5793(03)01062-7 (2003).

15. Boothby, T. C. & Pielak, G. J. Intrinsically Disordered Proteins and Desiccation Tolerance: Elucidating Functional and Mechanistic Underpinnings of Anhydrobiosis. Bioessays 39, (2017).

16. Tunnacliffe, A., Lapinski, J. & McGee, B. A Putative LEA Protein, but no Trehalose, is Present in Anhydrobiotic Bdelloid Rotifers. Hydrobiologia 546, 315–321 (2005).

17. Caprioli, M. et al. Trehalose in desiccated rotifers: a comparison between a bdelloid and a monogonont species. Comp. Biochem. Physiol. A Mol. Integr. Physiol. 139, 527–532 (2004).

18. Romero-Perez, P. S., Dorone, Y., Flores, E., Sukenik, S. & Boeynaems, S. When phased without water: Biophysics of cellular desiccation, from biomolecules to condensates. Chem. Rev. (2023) doi:10.1021/acs.chemrev.2c00659.

19. Tunnacliffe, A., Hincha, D. K., Leprince, O. & Macherel, D. LEA Proteins: Versatility of Form and Function. in Dormancy and Resistance in Harsh Environments (eds. Lubzens, E., Cerda, J. & Clark, M.) 91–108 (Springer Berlin Heidelberg, 2010).

20. Dure, L., Greenway, S. C. & Galau, G. A. Developmental biochemistry of cottonseed embryogenesis and germination: changing messenger ribonucleic acid populations as shown by in vitro and in vivo protein synthesis. Biochemistry 20, (1981).

21. Battista, J. R., Park, M. J. & McLemore, A. E. Inactivation of two homologues of proteins presumed to be involved in the desiccation tolerance of plants sensitizes Deinococcus radiodurans R1 to desiccation. Cryobiology 43, (2001).

22. Browne, J., Tunnacliffe, A. & Burnell, A. Plant desiccation gene found in a nematode. Nature 416, 38–38 (2002).

23. Schokraie, E. et al. Proteomic Analysis of Tardigrades: Towards a Better Understanding of Molecular Mechanisms by Anhydrobiotic Organisms. PLoS One 5, (2010).

24. Kikawada, T. et al. Dehydration-induced expression of LEA proteins in an anhydrobiotic chironomid. Biochem. Biophys. Res. Commun. 348, (2006).

25. Hand, S. C., Menze, M. A., Toner, M., Boswell, L. & Moore, D. LEA proteins during water stress: not just for plants anymore. Annu. Rev. Physiol. 73, (2011).

26. Battaglia, M., Olvera-Carrillo, Y., Garciarrubio, A., Campos, F. & Covarrubias, A. A. The Enigmatic LEA Proteins and Other Hydrophilins. Plant Physiol. 148, 6 (2008).

27. Yamaguchi, A. et al. Two Novel Heat-Soluble Protein Families Abundantly Expressed in an Anhydrobiotic Tardigrade. PLoS One 7, e44209 (2012).

28. Boothby, T. C. et al. Tardigrades Use Intrinsically Disordered Proteins to Survive Desiccation. Mol. Cell 65, 975 (2017).

29. Tanaka, S. et al. Novel Mitochondria-Targeted Heat-Soluble Proteins Identified in the Anhydrobiotic Tardigrade Improve Osmotic Tolerance of Human Cells. PLoS One 10, e0118272 (2015).

30. Garay-Arroyo, A., Colmenero-Flores, J. M., Garciarrubio, A. & Covarrubias, A. A. Highly Hydrophilic Proteins in Prokaryotes and Eukaryotes Are Common during Conditions of Water Deficit *. J. Biol. Chem. 275, 5668–5674 (2000).

31. Reyes, J. L. et al. Hydrophilins from distant organisms can protect enzymatic activities from water limitation effects in vitro. Plant Cell Environ. 28, 709–718 (2005).

32. Hesgrove, C. & Boothby, T. C. The biology of tardigrade disordered proteins in extreme stress tolerance. Cell Commun. Signal. 18, (2020).

33. Habchi, J., Tompa, P., Longhi, S. & Uversky, V. N. Introducing protein intrinsic disorder. Chem. Rev. 114, 6561–6588 (2014).

34. A mitochondrial late embryogenesis abundant protein stabilizes model membranes in the dry state. Biochimica et Biophysica Acta (BBA) - Biomembranes 1798, 1926–1933 (2010).

35. Cuevas-Velazquez, C. L., Saab-Rincón, G., Reyes, J. L. & Covarrubias, A. A. The Unstructured N-terminal Region of Arabidopsis Group 4 Late Embryogenesis Abundant (LEA) Proteins Is Required for Folding and for Chaperone-like Activity under Water Deficit *. J. Biol. Chem. 291, 10893–10903 (2016).

36. Bremer, A., Wolff, M., Thalhammer, A. & Hincha, D. K. Folding of intrinsically disordered plant LEA proteins is driven by glycerol-induced crowding and the presence of membranes. FEBS J. 284, (2017).

37. Hincha, D. K. & Thalhammer, A. LEA proteins: IDPs with versatile functions in cellular dehydration tolerance. Biochem. Soc. Trans. 40, (2012).

38. LeBlanc, B. M. & Hand, S. C. Target enzymes are stabilized by AfrLEA6 and a gain of α-helix coincides with protection by a group 3 LEA protein during incremental drying. Biochim. Biophys. Acta: Proteins Proteomics 1869, 140642 (2021).

39. Sowemimo, O. T. et al. Conserved Glycines Control Disorder and Function in the Cold-Regulated Protein, COR15A. Biomolecules 9, (2019).

40. Sanchez-Martinez, S., et al. Labile assembly of a tardigrade protein induces biostasis. *bioRxiv* 2023.06.30.547219 (2023) doi:10.1101/2023.06.30.547219.

41. Eicher, J. E. et al. Secondary structure and stability of a gel-forming tardigrade desiccation-tolerance protein. Protein Sci. 31, e4495 (2022).

42. Malki, A. et al. Intrinsically Disordered Tardigrade Proteins Self-Assemble into Fibrous Gels in Response to Environmental Stress. Angew. Chem. Int. Ed Engl. 61, (2022).

43. Piszkiewicz, S. et al. Protecting activity of desiccated enzymes. Protein Sci. 28, 941 (2019).

44. Packebush, M. H. et al. Natural and engineered mediators of desiccation tolerance stabilize Human Blood Clotting Factor VIII in a dry state. Sci. Rep. 13, 1–16 (2023).

45. Tanaka, A. et al. Stress-dependent cell stiffening by tardigrade tolerance proteins that reversibly form a filamentous network and gel. PLoS Biol. 20, e3001780 (2022).

46. Das, R. K. & Pappu, R. V. Conformations of intrinsically disordered proteins are influenced by linear sequence distributions of oppositely charged residues. Proc. Natl. Acad. Sci. U. S. A. 110, (2013).

47. Chandarlapaty, S. & Errede, B. Ash1, a daughter cell-specific protein, is required for pseudohyphal growth of Saccharomyces cerevisiae. Mol. Cell. Biol. 18, 2884– 2891 (1998).

48. Woolfson, D. N. & Williams, D. H. The influence of proline residues on α-helical structure. FEBS Lett. 277, 185–188 (1990).

50. 49. Deber, C. M., Brodsky, B. & Rath, A. Proline Residues in Proteins. in eLS (John Wiley & Sons, Ltd, 2010).

50. Holehouse, A. S., Das, R. K., Ahad, J. N., Richardson, M. O. G. & Pappu, R. V. CIDER: Resources to Analyze Sequence-Ensemble Relationships of Intrinsically Disordered Proteins. Biophys. J. 112, 16–21 (2017).

51. Batchelor, M. & Paci, E. Helical Polyampholyte Sequences Have Unique Thermodynamic Properties. J. Phys. Chem. B 122, 11784–11791 (2018).

52. Roesgaard, M. A. et al. Deciphering the Alphabet of Disorder—Glu and Asp Act Differently on Local but Not Global Properties. Biomolecules 12, 1426 (2022).

53. Martin, E. W. et al. Valence and patterning of aromatic residues determine the phase behavior of prion-like domains. Science 367, 694–699 (2020).

54. Mok, Y. K., Kay, C. M., Kay, L. E. & Forman-Kay, J. NOE data demonstrating a compact unfolded state for an SH3 domain under non-denaturing conditions. J. Mol. Biol. 289, 619–638 (1999).

55. Shimizu, T. et al. Desiccation-induced structuralization and glass formation of group 3 late embryogenesis abundant protein model peptides. Biochemistry 49, (2010).

56. Hundertmark, M., Popova, A. V., Rausch, S., Seckler, R. & Hincha, D. K. Influence of drying on the secondary structure of intrinsically disordered and globular proteins. Biochem. Biophys. Res. Commun. 417, (2012).

57. Shou, K. et al. Conformational selection of the intrinsically disordered plant stress protein COR15A in response to solution osmolarity – an X-ray and light scattering study. Phys. Chem. Chem. Phys. 21, 18727–18740 (2019).

58. Tunnacliffe, A. & Wise, M. J. The continuing conundrum of the LEA proteins. Naturwissenschaften 94, 791–812 (2007).

59. Yagi-Utsumi, M. et al. Desiccation-induced fibrous condensation of CAHS protein from an anhydrobiotic tardigrade. Sci. Rep. 11, 1–9 (2021).

60. Goyal, K., Walton, L. J. & Tunnacliffe, A. LEA proteins prevent protein aggregation due to water stress. Biochem. J 388, 151–157 (2005).

61. Chakrabortee, S. et al. Hydrophilic protein associated with desiccation tolerance exhibits broad protein stabilization function. Proc. Natl. Acad. Sci. U. S. A. 104, 18073 (2007).

62. Hatanaka, R., et al. An abundant LEA protein in the anhydrobiotic midge, PvLEA4, acts as a molecular shield by limiting growth of aggregating protein particles. (2013).

63. Bremer, A. et al. Intrinsically Disordered Stress Protein COR15A Resides at the Membrane Surface during Dehydration. Biophys. J. 113, (2017).

64. Miyazawa, K. et al. Tardigrade Secretory-Abundant Heat-Soluble Protein Has a Flexible β-Barrel Structure in Solution and Keeps This Structure in Dehydration. J. Phys. Chem. B 125, (2021).

65. Furuki, T. & Sakurai, M. Group 3 LEA protein model peptides protect liposomes during desiccation. Biochim. Biophys. Acta 1838, (2014).

66. Walker, J. M. The Proteomics Protocols Handbook. (Springer Science & Business Media, 2007).

67. Metapredict: a fast, accurate, and easy-to-use predictor of consensus disorder and structure. Biophys. J. 120, 4312–4319 (2021).

68. Mirdita, M. et al. ColabFold: making protein folding accessible to all. Nat. Methods 19, 679–682 (2022).

69. Pettersen, E. F. et al. UCSF ChimeraX: Structure visualization for researchers, educators, and developers. Protein Sci. 30, 70–82 (2021).

70. Eisenberg, D., Weiss, R. M. & Terwilliger, T. C. The helical hydrophobic moment: a measure of the amphiphilicity of a helix. Nature 299, 371–374 (1982).

71. Micsonai, A. et al. BeStSel: a web server for accurate protein secondary structure prediction and fold recognition from the circular dichroism spectra. Nucleic Acids Res. 46, W315–W322 (2018).

72. Drozdetskiy, A., Cole, C., Procter, J. & Barton, G. J. JPred4: a protein secondary structure prediction server. Nucleic Acids Res. 43, (2015).

