## Supplementary material for "Helicity of a tardigrade disordered protein promotes desiccation tolerance": File S1

CAHS Proteins from UniProt:

**>P0CU48.**

**CAHS 6_HYPEX**

**CAHS 94025**

MSGRNVESHMERNEKVVVNNSGHADVKKQQQQVEHTEFTHTEVKAPLIHPAPPIISTGAAGLAEEIVGQGFTASAARISGGTAEVHLQPSAAMTEEARRDQERYRQEQESIAKQQEREMEKKTEAYRKTAEAEAEKIRKELEKQHARDVEFRKDLIESTIDRQKREVDLEAKMAKRELDREGQLAKEALERSRLATNVEVNFDSAAGHTVSGGTTVSTSDKMEIKRN

**>P0CU47.**

**CAHS 5_HYPEX**

**CAHS 89226**

MATKESKYERVEKVNVDADGATLVKNIGEDRGKEDPGMNFQDKRPANLVPGAPAGVIPNRIESLPTDRAGQRLREHLSESERLRVSRSSTSSKSSSFVEPSLKYRGEIGPIGKNGEFVASSNRQNSSSNVSSSDNSERASPASRNSNPGMNNGMTTQRTTVITESSVQGLGAQRTVPIQPHQQREDHEVITHESHARAPETTVVTIPTTRFESAQLESRRDGRTYTEDKELTIPAPVVAPQIHAHQQVNMSGGTSATIHATTDLHLASEAQINDMGPEEYERYRAKVEALARIHEDETSRKAAAYRNAVEADAELIRQTLERQHMRDIEFRKDLVESSVDRQQQEIRLEAEYAMRALEQERVNARAALDQAMASTNIDVNIDSAIGTTHSQGRVTTTSESRTSQARGPATAAVI

**>P0CU46.**

**CAHS2_HYPEX**

**CAHS 86272** MSQQYEKKVERTEVVYGGDRRVEGSASASAEKTTNYTHTEIRAPMVNPLPPIISTGAAGLAQEIVGEGFTASATRISGAAATTQVLESQASREQAFKDQEKYSREQAAIARAHDKDLEKKTEEYRKTAEAEAEKIRKELEKQHARDVEFRKDLVESAIDRQKREVDLEAKYAKKELEHERELAMNALEQSKMATNVQVQMDTAAGTTVSGGTTVSEHTEVHDGKEKKSLGEKIKSLF

**>P0CU45.**

**CAHS1_HYPEX**

**CAHS 94025**

MSGRNVESHMERNEKVVVNNSGHADVKKQQQQVEHTEFTHTEVKAPLIHPAPPIISTGAAGLAEEIVGQGFTASAARISGGTAEVHLQPSAAMTEEARRDQERYRQEQESIAKQQEREMEKKTEAYRKTAEAEAEKIRKELEKQHARDVEFRKDLIESTIDRQKREVDLEAKMAKRELDREGQLAKEALERSRLATNVEVNFDSAAGHTVSGGTTISSSDKMEIKRN

**>P0CU51.**

**CAHS1_PARRC**

**CAHS 107838**

MSAEAMNMNMNQDAVFIPPPEGEQYERKEKQEIQQTSYLQSQVKVPLVNLPAPFFSTSFSAQEILGEGFQASISRISAVSEELSSIEIPELAEEARRDFAAKTREQEMLSANYQKEVERKTEAYRKQQEVEADKIRKELEKQHLRDVEFRKDIVEMAIENQKKMIDVESRYAKKDMDRERVKVRMMLEQQKFHSDIQVNLDSSAAGTETGGQVVSESQKFTERNRQIKQ

**>P0CU52.**

**CAHS2_PARRC**

**CAHS 106094**

MEAMNMNIPRDAMFVPPPESEQNGYHEKSEVQQTSYMQSQVKVPHYNFPTPYFTTSFSAQELLGEGFQASISRISAVTEDMQSMEIPEFVEEARRDYAAKTRENEMLGQQYEKELERKSEAYRKHQEVEADKIRKELEKQHMRDIEFRKEIAELAIENQKRMIDLECRYAKKDMDRERTKVRMMLEQQKFHSDIQVNLDSSAAGTESGGHVVSQSEKFTERNREMKR

**>P0CU43.**

**CAHS3_HYPEX**

**CAHS 77850**

MSNYQQESSYQYSDRSNNGQQQEQQEKKEVEHSSYTHTDVKVNMPNLIAPFISSSAGLAQELVGEGFQASVSRITGASGELTVIDTEAETEEARRDMEAKAREQELLSRQFEKELERKTEAYRKQQEVETEKIRKELEKQHLRDVEFRKELMEQTIENQKRQIDLEARYAKKELERERNKVKRVLERSKFHTDIQVNMEAAAGSTHSGSSSVAVSESEKFQTNN

**>P0CU44.**

**CAHS4_HYPEX**

**CAHS 77611**

MSNYQQESSYQYSDRSNNGQQQEQQEKKEVEHSSYTHTDVKVNMPNLIAPFISSSAGLAQELVGEGFQASVSRITGASGELTVIDTEAETEEARRDLEAKAREQELLSRQFEKELERKTEAYRKQQEVETEKIRKELEKQHLRDVEFRKELMEQTIENQKRQIDLEARYAKKELERERNKVKRVLERSKFHTDIQVNMEAAAGSTHSGSSSVAVSESEKFQTNN

**>J7M799.**

**CAHS1_RAMVA**

**CAHS1**

MPYEKHVEQTVVEKTEQPGHSSTHHAPAQRTVAREQEEVVHKEFTHTDIRVPHIDAPPPIIAASAVGLAEEIVSHGFQASAARISGASTEVDMRPSPKLAEEARRDAERYQKEHEMINRQAEATLQKKAEEYRHQTEAEAEKIRRELEKQHERDIQFRKDLIDQTIEKQKREVDLEAKMAKRELDREAQLAKEALERSRMATNVEVTLDTAAGHTVSGGTTVSSVDKVETVRERKHH

**>J7MDG6.**

**CAHS2_RAMVA**

**CAHS2** MSRDQGSTEYDANQRQEQHQEQHNTSYTHTDVRTNIPNIPAPFISTGVSGLGQQLVGEGFTASAARISGQSSETHVQMTPEMEAEARKDRERYERELQAINERHQRDIEGKTEAYRKQAEQEAERLRKELEKQHQRDIEFRKSLVQGTIENQKRQVELEAQLAKRELDREARLATQALDQSKMATDVQVNFDSAVGHTVSGATTVSQSEKVTQSKH

**>J7M3T1.**

**CAHS3_RAMVA**

**CAHS3**

MSSRQNQQSSSQHSSSSQQGGQGGQGVQGSSSYSRTEVHTSSGGPTIGGAQRTVPVPPGSHSEVHEEREVIKHGTKTESETHVVTVPVTTFGSTNMESVRTGFTVTQDKNLTVAAPNIAAPIHSNLDLNLGGGARAEITAGTTVDLSKIQRKDLGPEEYARYKAKVEQLARQDEQDAGMRAAQYREEVERDAELIRQILERQHIRDLEFRKEMVENQVNRQEREIQLEAEYAMRALELERNAAKEALESAKAQTNVNVKVESAIGTTVSKGAIQTSADKSSTTKTGPTTVTQIKHTEQHTERR

**>P0CU50.**

**CAHS8_HYPEX**

**CAHS 94063**

MSGRNVESHMERNEKVVVNNSGHADVKKQQQQVEHTEFTHTEVKAPLIHPAPPIISTGAAGLAEEIVGQGFTASAARISGGTAEVHLQPSAAMTEEARRDQERYRQEQESIAKQQEREMEKKTEAYRKTAEAEAEKIRKELEKQHARDVEFRKDLIESTIDRQKREVDLEAKMAKRELDREGQLAKEALERSRLATNVEVNFDSAAGHTVSGGTTVSTSDKMEIKRN


**CAHS D and its variants used in the study:**

**>CAHS D**

MSGRNVESHMERNEKVVVNNSGHADVKKQQQQVEHTEFTHTEVKAPLIHP

APPIISTGAAGLAEEIVGQGFTASAARISGGTAEVHLQPSAAMTEEARRD

QERYRQEQESIAKQQEREMEKKTEAYRKTAEAEAEKIRKELEKQHARDVE

FRKDLIESTIDRQKREVDLEAKMAKRELDREGQLAKEALERSRLATNVEV

NFDSAAGHTVSGGTTVSTSDKMEIKRN

**>N-terminus**

MSGRNVESHMERNEKVVVNNSGHADVKKQQQQVEHTEFTHTEVKAPLIHPAPPIISTGAAGLAEEIVGQGFTASAARISGGTAEVHLQPS

**>Linker region**

AAMTEEARRDQERYRQEQESIAKQQEREMEKKTEAYRKTAEAEAEKIRKELEKQHARDVEFRKDLIESTIDRQKREVDLEAKMAKRELDREGQLAKEALERSRLA

**>C-terminus**

TNVEVNFDSAAGHTVSGGTTVSTSDKMEIKRN

**>2X LR**

MSGRNVESHMERNEKVVVNNSGHADVKKQQQQVEHTEFTHTEVKAPLIHPAPPIISTGAAGLAEEIVGQGFTASAARISGGTAEVHLQPSAAMTEEARRDQERYRQEQESIAKQQEREMEKKTEAYRKTAEAEAEKIRKELEKQHARDVEFRKDLIESTIDRQKREVDLEAKMAKRELDREGQLAKEALERSRLATNMTEEARRDQERYRQEQESIAKQQEREMEKKTEAYRKTAEAEAEKIRKELEKQHARDVEFRKDLIESTIDRQKREVDLEAKMAKRELDREGQLAKEALERSRLATNVEVNFDSAAGHTVSGGTTVSTSDKMEIKRN

**>Ash1 LR**

MSGRNVESHMERNEKVVVNNSGHADVKKQQQQVEHTEFTHTEVKAPLIHPAPPIISTGAAGLAEEIVGQGFTASAARISGGTAEVHLQPSAAMTPSPSTPTKSGKMRSRSSSPVRPKAYTPSPRSPNYHRFALDSPPQSPRRSSNSSITKKGSRRSSGSSPTRPSPSTPTKSGKMRSRSSSPVRPKAYTPSPRLATNVEVNFDSAAGHTVSGGTTVSTSDKMEIKRN

**>LR_KToR**

AAMTEEARRDQERYRQEQESIARQQEREMERRTEAYRRTAEAEAERIRRELERQHARDVEFRRDLIESTIDRQRREVDLEARMARRELDREGQLAREALERSRLA

**>LR_RToK**

AAMTEEAKKDQEKYKQEQESIAKQQEKEMEKKTEAYKKTAEAEAEKIKKELEKQHAKDVEFKKDLIESTIDKQKKEVDLEAKMAKKELDKEGQLAKEALEKSKLA

**>LR_DToE**

AAMTEEARRDQERYRQEQESIAKQQEREMEKKTEAYRKTAEAEAEKIRKELEKQHAREVEFRKELIESTIERQKREVELEAKMAKRELEREGQLAKEALERSRLA

**>LR_EToD**

AAMTDDARRDQDRYRQDQDSIAKQQDRDMDKKTDAYRKTADADADKIRKDLDKQHARDVDFRKDLIDSTIDRQKRDVDLDAKMAKRDLDRDGQLAKDALDRSRLA

**>LR_P**

APMTEEPRRDPERYRPEQESIPKQQERPMEKKPEAYRKTPEEPEKIRKELEKPHARDVEFPRKDLIEPTIDRQKRPEVDLEPKMPKRELDREGQLPKEPLERSRL

**>LR_NH**

GGSTEEGRRDQERYRQEQESSGKQQERESEKKTEGYRKTGEGEGEKSRKESEKQHGRDSEFRKDSSESTSDRQKRESDSEGKSGKRESDREGQSGKEGSERSRSG

**>FL_LR_P**

MSGRNVESHMERNEKVVVNNSGHADVKKQQQQVEHTEFTHTEVKAPLIHPAPPIISTGAAGLAEEIVGQGFTASAARISGGTAEVHLQPSAPMTEEPRRDPERYRPEQESIPKQQERPMEKKPEAYRKTPEEPEKIRKELEKPHARDVEFPRKDLIEPTIDRQKRPEVDLEPKMPKRELDREGQLPKEPLERSRLTNVEVNFDSAAGHTVSGGTTVSTSDKMEIKRN

**>FL_LR_NH**

MSGRNVESHMERNEKVVVNNSGHADVKKQQQQVEHTEFTHTEVKAPLIHPAPPIISTGAAGLAEEIVGQGFTASAARISGGTAEVHLQPSGGSTEEGRRDQERYRQEQESSGKQQERESEKKTEGYRKTGEGEGEKSRKESEKQHGRDSEFRKDSSESTSDRQKRESDSEGKSGKRESDREGQSGKEGSERSRSGTNVEVNFDSAAGHTVSGGTTVSTSDKMEIKRN

**>Scramble_1**

MSGRNVESHMERNEKVVVNNSGHADVKKQQQQVEHTEFTHTEVKAPLIHPAPPIISTGAAGLAEEIVGQGFTASAARISGGTAEVHLQPESAEEMTREQRDAYRRQQESIEKAQEEMERQTKKAYETQKRAAKEIKELAERHLEAADDFVRRLEKISEDTIRVQKRLVEDMEKAAKRGLDLQEELAKEARESARKTNERVNFDSAAGHTVSGGTTVSTSDKMEIKRN

**>Scramble_2**

MSGRNVESHMERNEKVVVNNSGHADVKKQQQQVEHTEFTHTEVKAPLIHPAPPIISTGAAGLAEEIVGQGFTASAARISGGTAEVHLQPKSAMTEAEAQRYQRQSIDERAQEQMKERRTRARYTRQARDAEAIELHRKREKEFVEKLIARRKRSEDIRTEVEEEQDVELALMALKAKLGQLEEEDKAAKSETNVEKEDKNEFDSAAGHTVSGGTTVSTSDKMEIKRN
