## Supplementary material for "Helicity of a tardigrade disordered protein promotes desiccation tolerance": File S2

**R Script for CD spectra plots:**

library(readr)

library(dplyr)

library(ggplot2)

library(readxl)

library(ggsignif)

setwd("~/Downloads")

mre <- read_csv("xxx.csv")

cols = c("#2C8B57", "#609E9F", "#4C86C6", "#685EA9", "#7F469B")

plot(mre$wavelength, mre$'5', col = cols[1], type = "l", ylim = c(-4.5, 4), lwd = 3,

xlab = "Wavelength (nm)", ylab = "MRE", xlims = c(180, 270))

lines(mre$wavelength, mre$'5', col = cols[2], lwd = 3)

lines(mre$wavelength, mre$'12.5', col = cols[3], lwd = 3)

lines(mre$wavelength, mre$'25', col = cols[4], lwd = 3)

lines(mre$wavelength, mre$'50', col = cols[5], lwd = 3)

#cols= color codes different for different constructs

#mre$x= concentrations
